# Local jasmonic acid cues drive systemic acquired resistance signal generation

**DOI:** 10.1101/2021.01.20.427489

**Authors:** Jennifer Sales, Elisabeth Pabst, Marion Wenig, Heiko H. Breitenbach, Gerardo Perez, Claudia Knappe, Richard Hammerl, Jinghui Liu, Wilfried Rozhon, Brigitte Poppenberger, Erwin Grill, Corinna Dawid, A. Corina Vlot

## Abstract

The phytohormones salicylic acid (SA) and jasmonic acid (JA) promote two, mutually antagonistic immune pathways respectively protecting plants from biotrophic pathogens and necrotrophic pathogens or insects. This trade-off largely precludes the exploitation of SA and JA immune components for crop protection, raising the interest in immune signalling components that disrupt SA-JA antagonism. A local pathogen infection primes SA-dependent immunity in systemic tissues. This so-called systemic acquired resistance (SAR) ensures a long-lasting, broad-spectrum disease resistance that is not subject to SA-JA antagonism. Here, we show that two sequence-related LEGUME LECTIN-LIKE PROTEINs (LLPs) promote SAR through spatially separated functions with JA promoting local SAR signal generation through LLP3. In concert with LLP1, which is important for systemic recognition and propagation of SAR signals, LLP3 promotes both SA-dependent SAR and JA-mediated immunity. Thus, exploitation of LLP-associated signalling cues might allow application of plant innate immune signals to promote (crop) plant health.

## Introduction

As plants lack the dedicated immune cells and complex homeostatic systems that are found in animals, they developed alternative strategies of dealing with stress. An important aspect of this is the action of phytohormones and their associated signalling pathways. A key threat to plants comes from biotrophic or hemibiotrophic pathogens, which are fended off through salicylic acid (SA)-dependent responses induced at the site of infection. Pathogen-associated molecular patterns (PAMPs) from virulent pathogens are recognised by Pattern Recognition Receptors (PRRs) that are localised at the plasma membrane and initiate PAMP-triggered immunity (PTI)^1^. Effectors from avirulent pathogens are recognised by intracellular nucleotide-binding domain and leucine-rich repeat proteins (NLRs), and initiate the relatively stronger effector-triggered immunity (ETI)^2^. Both PTI and ETI responses rely on SA. *ENHANCED DISEASE SUSCEPTIBILITY1 (EDS1)* acts as a central regulator upstream of SA driving a positive feedback loop with SA to fortify defence^3,4^. Once local SA cascades are triggered, a systemic signal is generated to upregulate defence in distal plant parts. This broad spectrum response is known as Systemic Acquired Resistance (SAR). EDS1 is required for a successful SAR response, as evidenced by defects in both SAR signal generation and recognition in *eds1* mutant plants^5^.

During ETI, LEGUME LECTIN-LIKE PROTEIN1 (LLP1) accumulates in apoplast-enriched extracts of the model plant *Arabidopsis thaliana* in an EDS1-dependent manner^5^. *LLP1* is essential for SAR, primarily functioning in the systemic tissue in SAR signal perception or propagation^5,6^. LLP2 (At3g16530), which shares 66% similarity at the amino acid (AA) level with LLP1, was identified as a possible SAR-associated protein along with LLP1^5^. LLP1 and LLP2 respectively share 61% and 87% AA similarity with LLP3 (At3g15356; LECTIN in^7^). LLP2 and LLP3 are induced at the transcriptional level by chitin and jasmonic acid (JA), respectively^7,8^, but their physiological roles remain unknown.

For a functional SAR response to occur, two interconnected signalling pathways are required^9,10^. The first of these pathways is primarily associated with SA^4,11^. The second involves pipecolic acid (Pip), and its presumed bioactive derivative *N-* hydroxy-pipecolic acid (NHP)^12–14^. LLP1 has a key role in the latter cascade acting downstream of Pip and upstream of SAR-associated volatile signals to propagate SAR-associated immunity in systemic tissues^6^.

When considering plants in complex natural systems, biotrophic defence signalling cascades associated with SA interact with other stress response pathways. These include abiotic stress responses associated with abscisic acid (ABA) and defence against necrotrophic pathogens and insects controlled by JA^15–17^. In order to fine tune defence, and optimise resource allocation, there is general antagonism between the three pathways^18^. Studies in Arabidopsis, for example, showed that after SA defences were activated by a hemibiotrophic pathogen, plants also became more susceptible to a necrotrophic pathogen, *Alternaria brassicicola,* pointing towards a down-regulation in JA-mediated defences^19,20^. Interestingly, the same studies found that this antagonism was restricted to the infected tissues and did not spread systemically during SAR. Recent evidence further convolutes the role of JA in SA-mediated defence which appears to be highly dependent on concentration, spatial distribution, and circadian rhythm^21–23^.

It is common for plants to be challenged by abiotic factors at the same time that they are under pathogen attack. Indeed, as climates shift, not only are traditional crops placed under greater stress from factors such as drought and salinity, but also from emerging infectious diseases and existing pathogens that have expanded their geographical range^24,25^. This threat to crop security increases the necessity for knowledge of interactions between the different stress pathways. Here we show that *LLP1* and *LLP3* have novel functions in multiple stress responses, and harbor significant potential for engineering multi-stress tolerance in plants.

## Results

### LLP3 is essential for local SAR signal generation

SA is a key component for plant defence against biotrophic pathogens, in both local and systemic tissues. Here, both Col-0 wild type and *eds1-2* knockout plants were spray-treated with 1 mM SA. After 24 h, the levels of *LLP1* transcripts were induced, and this effect was not dependent on EDS1 (Fig. 1A;^5^). In contrast to Lyou et al.^7^, who detected a slight reduction in *LLP3* transcript abundance after treatment of plants with 50 μM SA, we did not observe a reproducible down regulation of *LLP3.* Similarly, *LLP2* transcript levels did not significantly change in response to SA in either genotype. Similar results were seen if plants were treated with the SA analogue 1,2,3-benzothiadiazole-7-carbothioic acid *S*-methyl ester (BTH; Supplementary Fig. 1). Thus, *LLP1* transcript accumulation, and not that of its homologues *LLP2* and *LLP3*, is regulated by SA, and this regulation is independent of EDS1.

**Figure 1.**
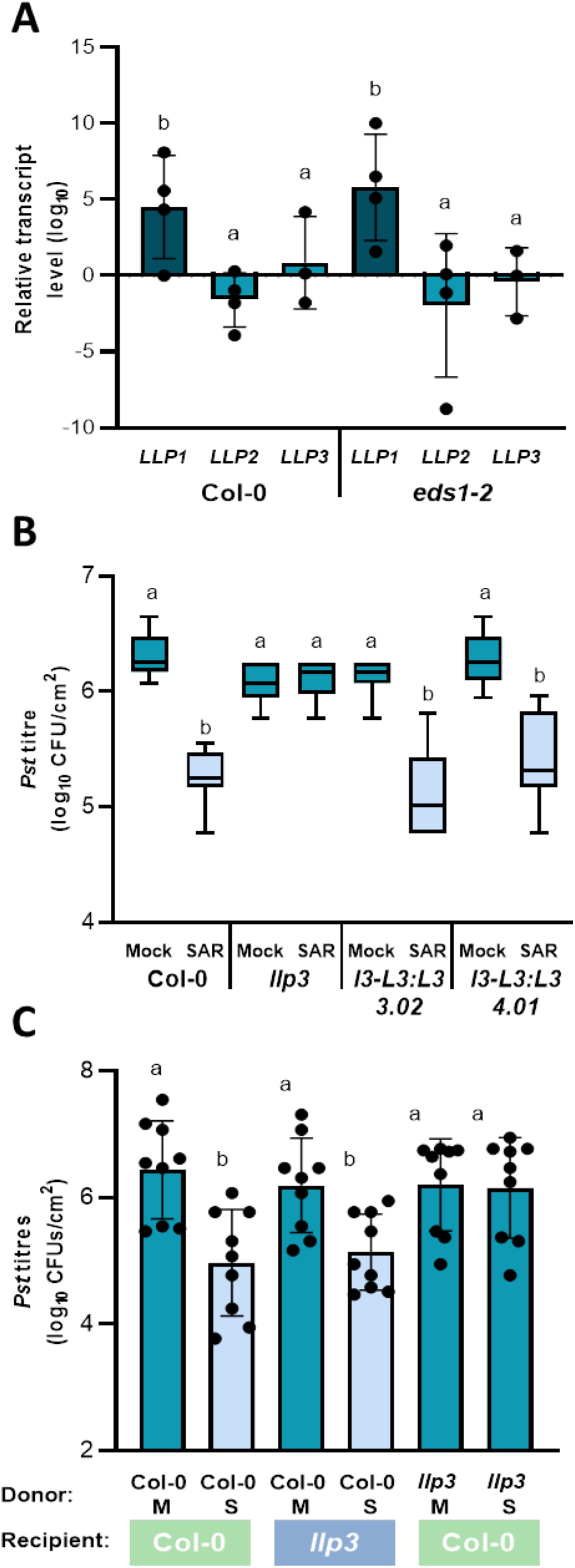
*LEGUME LECTIN-LIKE PROTEIN3 (LLP3)* promotes systemic acquired resistance (SAR) signal generation/transmission. (A) *LLP3* transcript accumulation is not affected by salicylic acid (SA). 4-week-old Col-0 wild type and *eds1-2* Arabidopsis plants were spray-treated with 1mM SA, and 24 hours (h) later *LLP1, LLP2,* and *LLP3* transcript accumulation was determined by RT-qPCR. Transcript accumulation was normalized to that of *UBIQUITIN* and is shown relative to the normalized transcript levels in the appropriate mock controls. Bars represent the log_2_(mean) ± SEM of four biologically independent replicates. The letters above the bars indicate statistically significant differences (one-way ANOVA, n=4, P<0.05, F=4.623, DF=22). (B) *LLP3* is required for SAR. Plants were infiltrated locally with either *Pst AvrRpm1* (SAR) or with 10 mM MgCl2 as the mock control (M). To monitor SAR, 3 days after the primary treatment leaves distal to the initial treatment site were infiltrated with *Pst*. Plant lines included *llp3* mutants and 2 independently transformed complementation lines carrying a transgene driving *LLP3* expression from its native promoter (*l3-L3:L3* 3.02 and 4.01). Box plots represent average *Pst* titres in systemic leaves at 4 days post-inoculation (dpi) from 4 biologically independent experiments, including 3 replicates each ± min and max values. The letters above the box plots indicate statistically significant differences (Kruskal-Wallis test, P<0.05, n=12, KW statistic=101.4). (C) *LLP3* is required to send, but not to receive phloem-mobile SAR signals. Leaves of donor plants were inoculated with *Pst AvrRpm1* (S) or with the appropriate mock control (M). After 24 h, petiole exudates were collected from the donor plants and infiltrated into leaves of naïve recipient plants. 24h later, the treated leaves were challenged with *Pst*. Bars represent average *Pst* titres at 4 dpi from 3 biologically independent experiments, including 3 replicates each ± SD. The letters above the bars indicate statistically significant differences (one-way ANOVA, n=9, P<0.05, F=6.258, DF=35).

Reduced transcript accumulation of *LLP1*, *LLP2*, and *LLP3* in *RNAi:LLP1-3* plants compromises the ability of the plants to generate or transmit phloem-mobile SAR signal(s)^6^. Because we have so far been unable to generate viable *llp2* mutant plants, we focused on *LLP3*, whose transcript accumulation was reduced in an *llp3* T-DNA mutant (Supplementary Fig. 2). To analyze SAR, these plants were initially infiltrated in two leaves with either *Pst AvrRpm1* or a 10 mM MgCl2 mock control solution. Three days later, two leaves distal to the initial infection were infiltrated with virulent *Pst*. After another four days, the resulting *in planta Pst* titres were determined. In wildtype plants, a local *Pst AvrRpm1* infection reduced *Pst* growth in the systemic tissues compared to that in mock-treated plants, indicating the establishment of SAR (Fig. 1B). SAR was fully abolished in *llp3* mutant plants (Fig. 1B). Ectopic expression of a wildtype copy of *LLP3* driven by its native promoter in the *llp3* mutant background (*llp3-LLP3:LLP3*) raised *LLP3* transcript accumulation to intermediate levels between that of wildtype and *llp3* mutant plants (Supplementary Fig. 2). This complemented the SAR-defective phenotype of the *llp3* mutant (Fig. 1B), ascribing LLP3 an essential role in SAR.

We next tested if *LLP3* acts locally or systemically in SAR by using petiole exudates (PEX) from *Pst AvrRpm1-*inoculated and mock-treated plants. 24 Hours (h) after infiltration of these PEX in recipient plants, the treated leaves were inoculated with *Pst* and the resulting *Pst* titers monitored at 4 days post-inoculation (dpi). PEX from infected wildtype plants reduced *Pst* growth in wildtype recipient plants as compared to PEX from mock-treated wildtype plants (Fig. 1C). Similarly, *llp3* recipient plants responded with reduced *Pst* growth to PEX from infected wildtype plants, suggesting that *LLP3* is not involved in systemic recognition or propagation of SAR signal(s). In contrast, PEX from infected *llp3* donor plants did not reduce *Pst* growth in wildtype recipient plants (Fig. 1C), suggesting that *LLP3* is necessary for local SAR signal generation or transmission. This spatially separates the role in SAR of *LLP3* from that of its sequence-related homolog *LLP1,* which acts systemically in SAR^6^.

### LLP1-3 influence responses to abiotic stress

Because *LLP3* did not show a significant response to SA treatment, but nevertheless influenced SAR, we questioned if *LLP3* might be regulated by phytohormones other than SA. Yasuda et al.^26^ showed that ABA and ABA-dependent responses to salinity stress compromised SA signalling and potentially SAR. In order to investigate whether ABA had an impact upon the transcript levels of *LLP1, LLP2,* and *LLP3,* plants were spray-treated with 100μM ABA and tissues were harvested 24 h later. In both Col-0 wild type and *eds1-2* plants, *LLP1* transcript levels were significantly downregulated, while there was no change in transcript levels of either *LLP2* or *LLP3* (Figure 2A). Thus, ABA downregulates *LLP1* transcript accumulation independently of *EDS1.*

**Figure 2.**
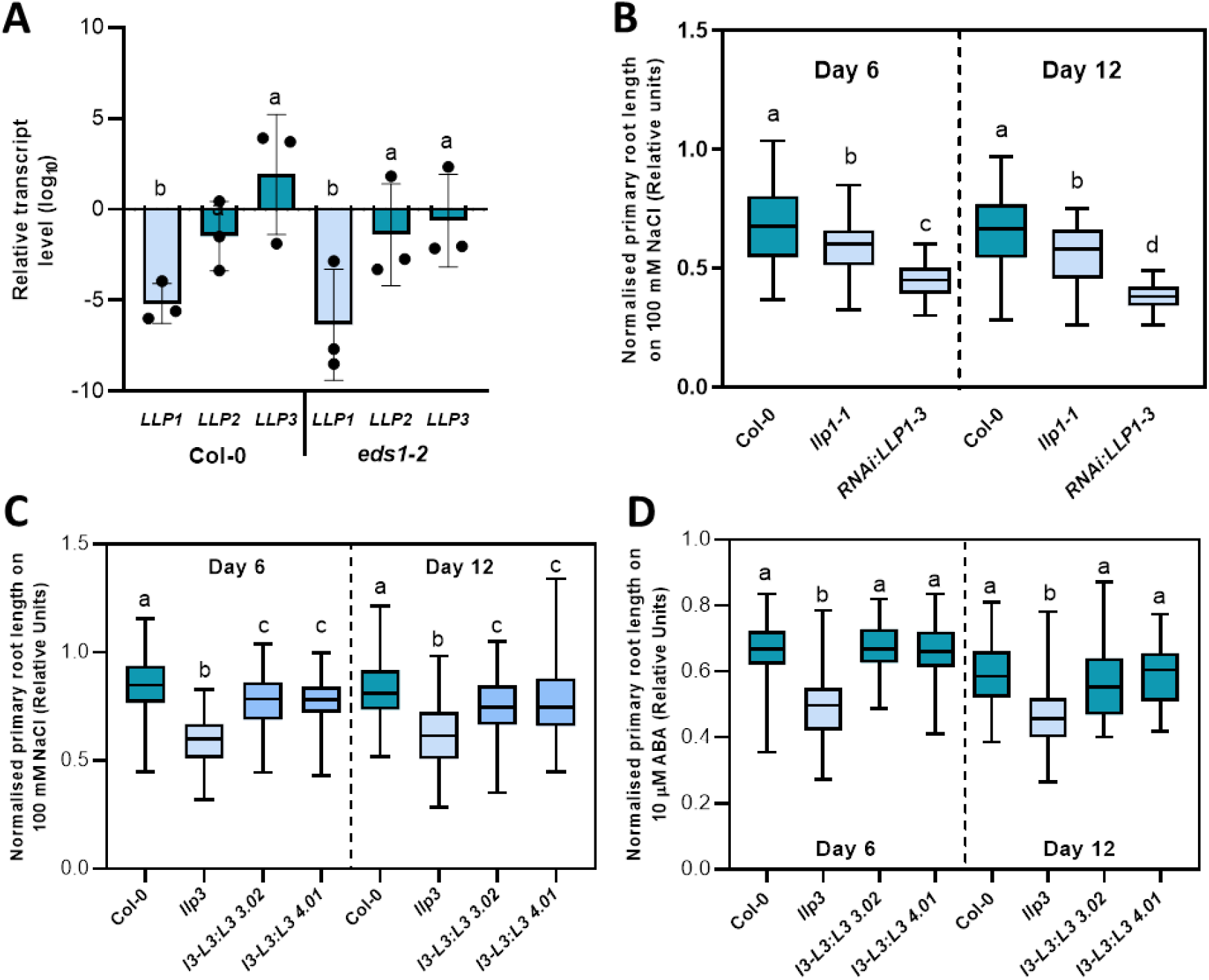
*LLP1, LLP2,* and/or *LLP3* compromise Arabidopsis responses to salt stress. (A) *LLP1* transcript accumulation is reduced after ABA treatment. Col-0 wild type and *eds1-2* plants were spray-treated with 100μM ABA, and after 24 h *LLP1, LLP2,* and/or *LLP3* transcript accumulation was determined by RT-qPCR. Transcript accumulation was normalized to that of *UBIQUITIN* and is shown relative to the normalized transcript levels in the appropriate mock controls. Bars represent the log_2_(mean) ± SEM of three biologically independent replicates. The letters above the bars indicate statistically significant differences (one-way ANOVA, n=3, P<0.05, F=6.291, DF=40). (B/C) *LLP1, LLP2,* and/or *LLP3* compromise salt-associated root growth inhibition. Seedlings of Col-0 wild type, *llp1-1,* and *RNAi:LLP1-3* (B) and of *llp3* and two *llp3-LLP3:LLP3* complementation lines *(l3-L3:L3* 3.02 and 4.01; C) were germinated on control MS plates, and after 6 days transferred to either further control plates, or to plates supplemented with 100 mM NaCl. Primary root length was measured at 6 and 12 days post transfer and normalized to that of the same genotype on control plates. Box plots represent average normalized root length ± min and max values. The letters above the box plots indicate statistically significant differences (B: one-way ANOVA, P=<0.05, F=30.70, DF=233, for day 6, Col-0 n=48, *llp1-1* n=38, *RNAi:LLP1* n=40, for day 12 Col-0 n=48, *llp1-1* n=29, *RNAi:LLP1-3* n=21; C: Day 6: Kruskal-Wallis test, P=<0.05, KW test statistic =165.5, Col-0 n=83, *llp3* n=86, *l3-L3:L3 3.02* n=96, *l3-L3:L3 4.01* n=89. Day 12: one-way ANOVA, P=<0.05, F=25.08, DF=519, Col-0 n=81, *llp3* n=84, *l3-L3:L3 3.02* n=94, *l3-L3:L3 4.01* n=84). These experiments were repeated 3 (C) to 4-8 times (B) with comparable results. (D) *LLP3* compromises root growth inhibition on 10 μM ABA. Col-0 wild type, *llp3* and two *llp3-LLP3:LLP3* complementation lines were treated as described in (B/C), and the treatment plates were supplemented with 10 μM ABA. Box plots represent average normalized root length ± min and max values. The letters above the box plots indicate statistically significant differences (Day 6: one-way ANOVA, P=<0.05, F=76.10, DF=538, Col-0 n=76, *llp3* n=86, 3.01 n=89, *l3-L3:L3 3.02* n=98, *l3-L3:L3 4.01* n=94, 8.01 n=96. Day 12: Kruskal-Wallis test, P=<0.05, KW test statistic =121.1, Col-0 n=85, *llp3* n=90, *l3-L3:L3 3.02* n=100, *l3-L3:L3 4.01* n=97). This experiment was repeated 3 times with comparable results.

ABA is an important phytohormone in abiotic stress signalling. Therefore, it seemed possible that if *LLP1* was transcriptionally regulated by ABA, *llp1-1* mutant and *RNAi:LLP1-3* plants may show an altered phenotype under abiotic stress. To test for aberrant reactions to high salinity, seedlings were germinated and after 6 days transferred to treatment plates with 100mM NaCl. The length of the primary roots was measured at 6 and 12 days post transfer and normalised to those on control plates (to which seedlings had also undergone transfer). While *llp1-1* plants had marginally longer roots than wildtype on control plates (Supplementary Fig. 3), the plants of all genotypes showed a significant reduction in root length when grown on salt compared to control conditions. Notably, both *llp1-1* and *RNAi:LLP1-3* plants showed more pronounced salt-induced root growth inhibition as compared to wildtype plants, and *RNAi:llp1-3* was significantly more affected than *llp1-1* (Fig. 2B).

To test for a possible contribution of *LLP3* to salt-induced root shortening, we transferred *llp3* seedlings and those of two *llp3-LLP3:LLP3* complementation lines to treatment plates with 100 mM NaCl. Similar to *llp1-1, llp3* mutants displayed exaggerated root growth inhibition under salt stress, and this phenotype was complemented by ectopic expression of *LLP3:LLP3* (Fig. 2C). This might be associated with changes in ABA responses, because ABA-induced root shortening was also exaggerated in the *llp3* mutant, but not in *llp3-LLP3:LLP3* complementation lines (Fig. 2D). However, ABA-induced root shortening was only moderately changed in *llp1-1* and not changed in *RNAi:LLP1-3* plants compared to wild type (Supplementary Fig. 4A). Also, ABA-induced transcript accumulation of the ABA marker gene *RAB18* was the same in all genotypes (Supplementary Fig. 4B/C). Therefore, a contribution of ABA to *LLP-*associated root shortening is probably minor. The absence of ABA-associated phenotypes was further supported by the response of *RNAi:LLP1-3* plants to progressive drought. There was no physiologically relevant or significant difference between *RNAi:LLP1-3* and wild type plants when water consumption and water use efficiency (WUE) under progressive drought, were examined (Supplementary Fig. 5). We therefore posit that the increased sensitivity of the *RNAi:LLP1-3* lines to high salinity is most likely mechanistically independent of ABA.

### LLP3 responds to MeJA and affects JA-mediated responses

Another candidate pathway that has been shown to be involved in both biotic defence and salt stress tolerance is the JA pathway^27^. Also, MeJA treatment has been associated with increased *LLP3* transcript levels^7^. Here, spray treatment of Arabidopsis with 100 μM MeJA increased accumulation of *LLP3* transcripts in both Col-0 and *eds1-2* plants, suggesting that MeJA induces *LLP3* in an *EDS1-* independent manner (Fig. 3A). The accumulation of *LLP1* and *LLP2* transcripts was not significantly changed by MeJA. We next investigated whether the reduction of LLP3 in *RNAi:LLP1-3* plants would affect JA-mediated defence against a necrotrophic pathogen. Indeed, lesions induced by *A. brassicicola*, a necrotrophic fungus, were larger in *RNAi:LLP1-3* plants than in wildtype plants and similar phenotypes were observed in *llp1-1* and *llp3* mutant plants (Fig. 3B/C). This shows that LLP1, LLP2, and/or LLP3 promote defence against necrotrophic pathogens and thus potentially normal JA signalling under biotic stress.

**Figure 3.**
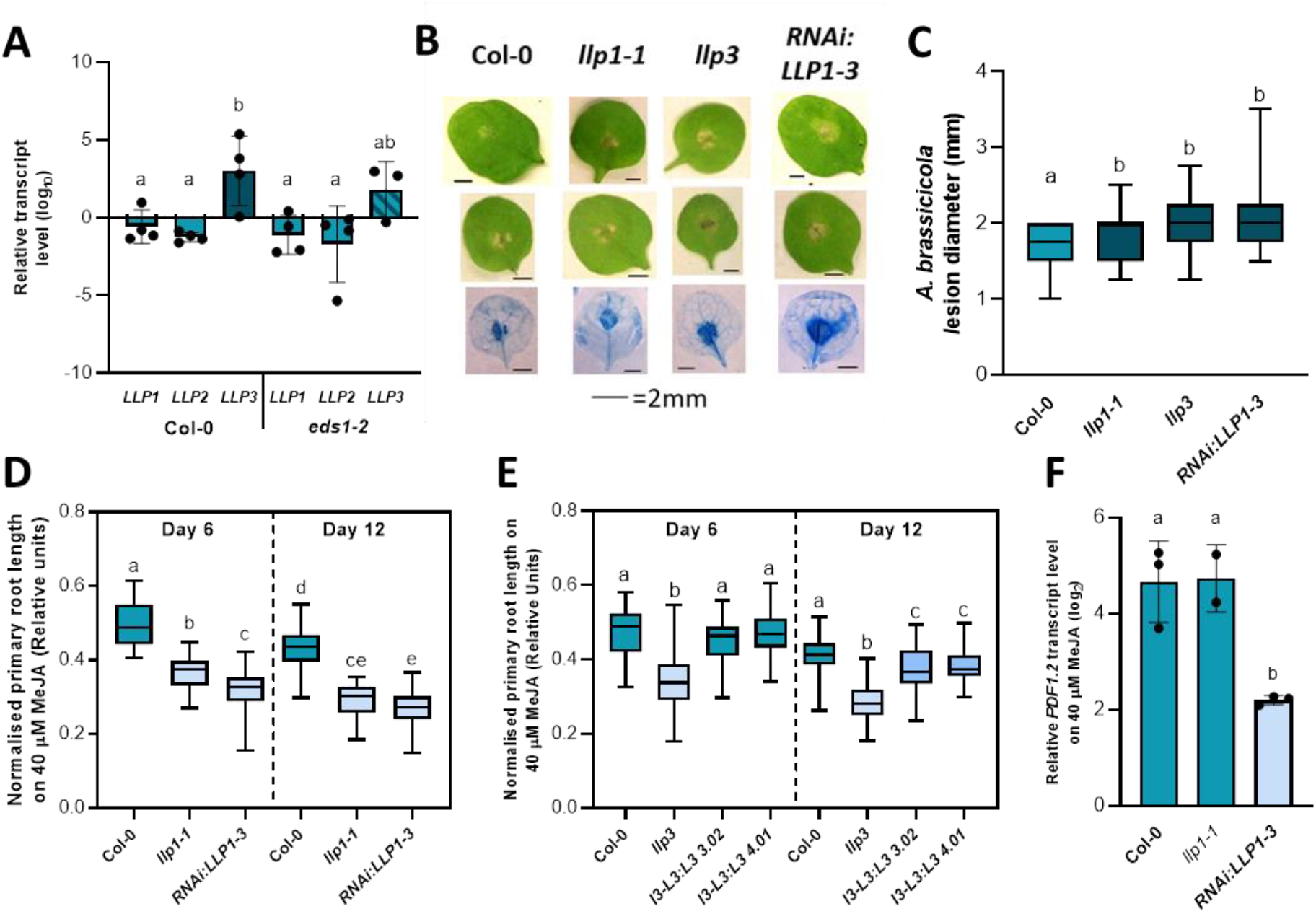
*LLP1, LLP2,* and/or *LLP3* differentially affect jasmonic acid (JA)-associated responses in Arabidopsis. (A) *LLP3* transcript accumulation is induced by methyl jasmonate (MeJA). Col-0 wild type and *eds1-2* plants were spray-treated with 100 μM MeJA, and after 24 h *LLP1, LLP2,* and/or *LLP3* transcript accumulation was determined by RT-qPCR. Transcript accumulation was normalized to that of *UBIQUITIN* and is shown relative to the normalized transcript levels in the appropriate mock controls. Bars represent the log2(mean) ± SEM of four biologically independent replicates. The letters above the bars indicate statistically significant differences (one-way ANOVA, n=4, P<0.05, F=4.493, DF=45). (B/C) *LLP1, LLP2,* and/or *LLP3* promote JA-associated defence against *Alternaria brassicicola.* Droplets containing spores of the necrotrophic fungus *A. brassicicola* were placed on the leaves of four-week-old Col-0 wild type, *llp1-1, llp3,* and *RNAi:LLP1-3* plants. Resulting lesions were photographed (B) and measured (C) 5 days later. Box plots in (C) represent mean lesion diameters from 4 biologically independent experiments including 15 replicates each ± min and max values. The letters above the box plots indicate statistically significant differences (Kruskal-Wallis test, P=<0.05, KW test statistic =24.10, n=60 for all genotypes). (D/E) *LLP1, LLP2,* and/or *LLP3* compromise JA-associated root growth inhibition. Seedlings of Col-0 wild type, *llp1-1,* and *RNAi:LLP1-3* (D) and of *llp3* and two *llp3-LLP3:LLP3* complementation lines (*l3-L3:L3* 3.02 and 4.01; E) were germinated on control MS plates, and after 6 days transferred to either further control plates, or to plates supplemented with 40 μM MeJA. Primary root length was measured at 6 and 12 days post transfer and normalized to that of the same genotype on control plates. Box plots represent average normalized root length ± min and max values. The letters above the box plots indicate statistically significant differences (D: Day 6: one-way ANOVA, F=44.87, DF=147, Col-0 n=29, *llp1-1, RNAi:LLP1-3* n=30. Day 12: one-way ANOVA, F=74.62, DF=175, Col-0 n=29, *llp1-1* n=30, *RNAi:LLP1-3 n=28;* E: Day 6: one-way ANOVA, P<0.05, F=61.40, DF=541, Col-0 n=71, *llp3* n=85, *l3-L3:L3 3.02* n=94, *l3-L3:L3 4.01* n=97. Day 12: Kruskal-Wallis test, P=<0.05, KW test statistic =140.7, Col-0 n=63, *llp3* n=73, *13-L3:L3 3.02* n=94, *l3-L3:L3 4.01* n=97, 8.01 n=95). These experiments were repeated 3 (E) to 5 times (D) with comparable results. (F) *LLP1, LLP2,* and/or *LLP3* compromise MeJA-induced *PDF1.2* transcript accumulation. *PDF1.2* transcript accumulation was monitored by qRT-PCR in seedlings from (D). Transcript accumulation was normalized to that of *UBIQUITIN* and is shown relative to the normalized transcript levels in the appropriate mock controls. Bars represent the log2(mean) ± SEM of biologically independent replicates. The letters above the bars indicate statistically significant differences (one-way ANOVA, P=<0.05, F=14.93, DF=7, n=3 for Col-0 and *RNAi:LLP1-3,* n=2 for *llp1-1).*

We subsequently tested if compromised JA signalling could have been responsible for the root growth inhibition phenotype of the *RNAi:LLP1-3* seedlings on salt. To this end, we again used the root growth inhibition assay, but this time the treatment plates were supplemented with 40 μM MeJA. This treatment induced similar results as treatment with NaCl. Both the *llp1-1* mutant and *RNAi:LLP1-3* seedlings showed significantly enhanced root length inhibition compared to wildtype (Fig. 3D). Similarly, *llp3* mutant seedlings displayed exaggerated root growth inhibition on MeJA and this phenotype was complemented by ectopic expression of *LLP3:LLP3* (Fig. 3E). JA downstream signalling pathways in the seedlings after 12 days on MeJA-supplemented plates were also aberrant in the *RNAi:LLP1-3* seedlings. Transcript accumulation of the JA marker gene *PDF1.2* was increased 12 days after transfer of wildtype, *llp1-1,* and *llp3* seedlings from control to MeJA plates (Fig. 3F and Supplementary Fig. 6A). By contrast, the induction of *PDF1.2* transcript accumulation was compromised in *RNAi:LLP1-3* plants (Fig. 3F), while the transcript accumulation of *VSP2* remained unchanged (Supplementary Fig. 6B).

To determine whether the loss of *LLP1-3* was affecting gene expression through changes in hormone biosynthesis, or through downstream interactions, the net content of SA, JA and ABA was measured in *RNAi:LLP1-3* plants after treatment with salt. Although there was an increase in net JA and ABA content after salt treatment, there was no significant difference between wildtype and *RNAi:LLP1-3* plants (Supplementary Fig. 7). This indicates that signalling aberration does not occur in the biosynthesis of these phytohormones, and that any crosstalk occurs in the pathways downstream of phytohormone biosynthesis. Thus, the three LLP proteins appear to simultaneously promote JA-associated defence against necrotrophic *A. brassicicola* and JA-associated salt tolerance in a process that is occurring downstream of JA accumulation.

### Crosstalk between JA and SA signalling pathways is misregulated in RNAi:LLP1-3 plants

From the above experiments, the *RNAi:LLP1-3* plants show a different level of JA marker gene transcript accumulation under abiotic stress. The SA and JA signalling pathways have multiple points of interaction, normally resulting in antagonistic cross talk1^8^. We therefore investigated *PR1* transcript accumulation as a marker of SA signalling after watering of mature plants with salt. Whereas *PR1* transcript levels were reduced in Col-0 wild type after salt treatment when compared to a mock control, possibly due to the antagonistic relationship between SA and either ABA or MeJA, *PR1* transcript levels remained unchanged in *llp1-1* and were upregulated by ~80-fold in salt-compared to mock-treated *RNAi:LLP1-3* plants (Fig. 4A). Hence, LLP1, LLP2, and/or LLP3 might co-operate in compromising responses to salt or associated JA-SA crosstalk events resulting in enhanced SA-associated responses in *RNAi:LLP1-3* plants in response to salinity stress.

**Figure 4.**
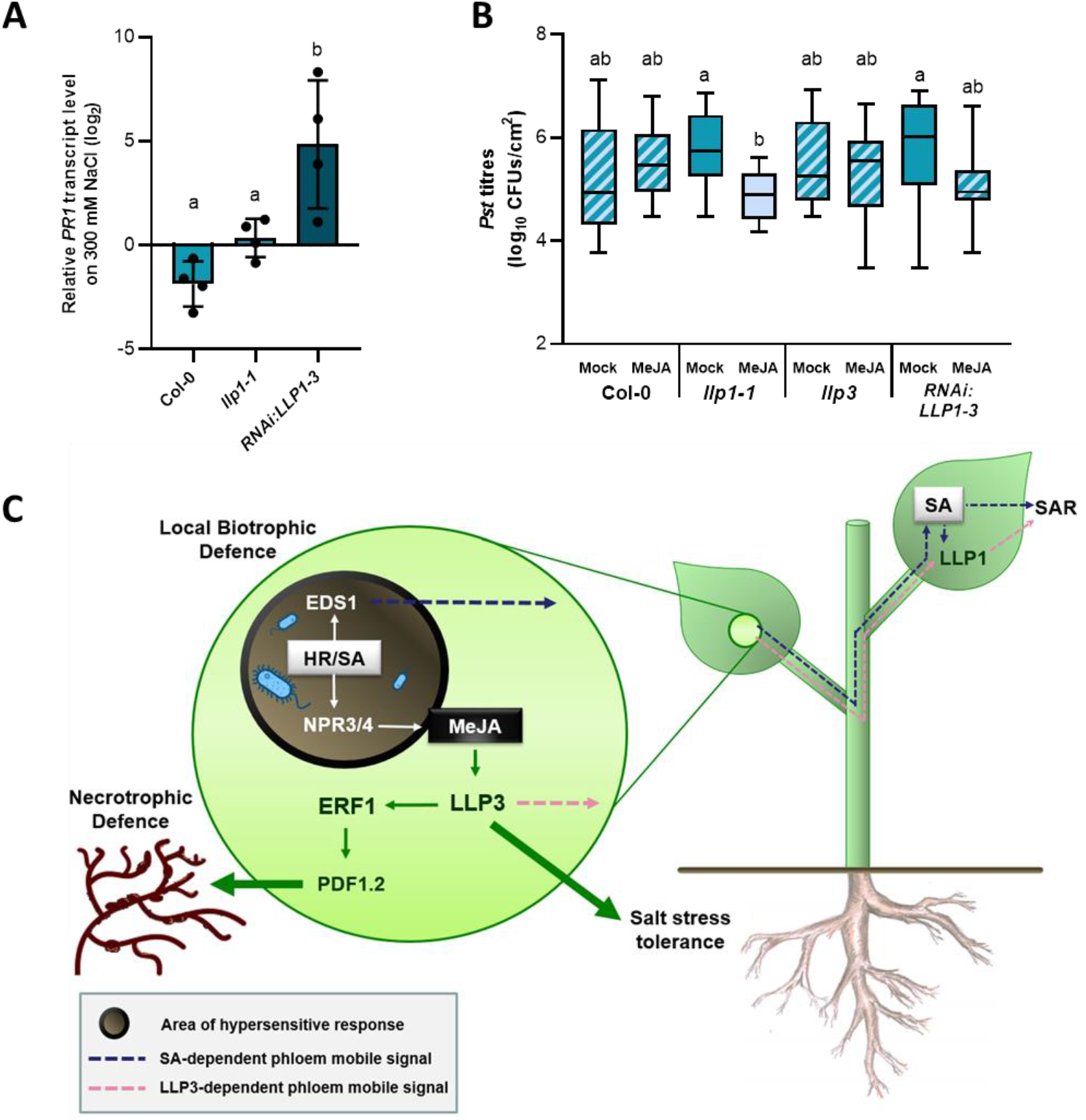
*LLP1, LLP2,* and/or *LLP3* dampen antagonistic SA-JA cross talk between defence pathways. (A) Exposure to salt drives up *PR1* transcript levels when *LLP1, LLP2,* and/or *LLP3* transcript levels are reduced. Four-week-old Col-0 wild type, *llp1-1,* and *RNAi:LLP1-3* plants were irrigated with 300 mM NaCl three times over the course of 9 days. Three days later, *PR1* transcript accumulation in the leaves was determined by qRT-PCR. Transcript accumulation was normalized to that of *UBIQUITIN* and is shown relative to the normalized transcript levels in the appropriate mock controls. Bars represent the log2(mean) ± SEM of four biologically independent replicates. The letters above the bars indicate statistically significant differences (one-way ANOVA, n=4, P<0.05, F=12.23, DF=11). (B) In the absence of functional *LLP1,* MeJA triggers SAR-like resistance in distal tissues. Col-0 wild type, *llp1-1, llp3,* and *RNAi:LLP1-3* plants were treated locally with 100 μM MeJA by leaf infiltration. To monitor systemic SA-associated defence responses, leaves distal to the site of the initial treatment were inoculated with *Pst* 3 days after the primary treatment. Box plots represent average *Pst* titres in systemic leaves at 4 dpi from 4-5 biologically independent experiments, including 3-4 replicates each ± min and max values. The letters above the box plots indicate statistically significant differences (Kruskal-Wallis test, P<0.05, KW test statistic=20.61, Col-0 mock n=17, MeJA n=20, *llp1-1* mock n=18, MeJA n=19, *llp3* mock n=11, MeJA n=11, *RNAi:LLP1-3* mock n=19, MeJA n=19). (C) *LLP3* promotes local SAR signal generation downstream of (Me)JA accumulating in the perimeter of HR (hypersensitive response) lesions. Elevated *LLP3* expression promotes *PDF1.2* expression and defence against necrotrophic pathogens through ERF1 (ETHYLENE RESPONSE FACTOR 1) as well as salt stress tolerance. In parallel with EDS1-dependent, SA-associated long distance signals, *LLP3* promotes accumulation or transmission of a long distance SAR signal downstream of (Me)JA. Systemically, *LLP1* balances incoming signals promoting SAR while restricting deleterious effects of SA-associated SAR on JA-associated defence responses. Abbreviations: NPR3/4, NON-EXPRESSOR OF PR GENES3/4

As MeJA-associated stress was able to induce SA-dependent gene expression in *LLP1-3*-compromised plants, we investigated whether a local application of MeJA would be sufficient to reconstitute a systemic defence response in the same lines. Using a similar experimental setup to a classical SAR assay, two lower leaves were infiltrated with 100 uM MeJA, and systemic leaves were infiltrated with virulent *Pst* three days later. The bacterial titres in the systemic leaves at 4 dpi indicated that MeJA, while not affecting bacterial titres in wildtype, was able to reconstitute a SAR response in *llp1-1* mutant plants, but not in *llp3* or *RNAi:LLP1-3* lines (Fig. 4B).

## Discussion

In this paper we show that LLP3 acts locally in SAR signal generation. *LLP3* expression is induced by MeJA, and *llp3* mutant plants display JA-associated biotic and abiotic stress tolerance phenotypes (Fig. 3). This implies that local JA responses contribute to SAR signal generation or transmission. Until now, a potential role of JA in SAR has been under debate^28,29^, and JA has been believed to be subject to antagonistic control in local infected tissues undergoing ETI1^9,20^. However, during RPS2-mediated ETI the accumulation of SA and downstream signalling through the NPR3 and NPR4 receptors initiates *de novo* JA synthesis^30^, and the SA sector in

Arabidopsis immune networks activated during PTI is dependent upon the JA sector^31^. Given that PTI and ETI have some convergent signalling pathways, including through SA accumulation^32,33^, it is likely that JA may have a more important role in biotrophic immunity than has been traditionally recognized.

SA and Pip are thought to function via interconnected signalling pathways during SAR^9,10^. *LLP1* transcript accumulation is increased in response to SA (Fig. 1A), but is dispensable for SA-induced immunity^5^. LLP1 further promotes systemic SAR signal recognition or propagation downstream of Pip and drives a positive feedback loop propagating volatile monoterpene emissions as airborne SAR cues^6^. Notably, MeJA-induced root growth inhibition was only marginally, if at all, exaggerated in Pip-deficient *ald1*^34^ plants and in different monoterpene emission-compromised^*6*^ mutants (Supplementary Fig. 8). Therefore, *LLP1* might interact with an *LLP3-*associated SAR signalling component in a pathway that is mostly separate from its role in Pip signalling and monoterpene transmission.

JA activates two separate signalling pathways, depending upon which other signals/factors are detected at the same time. This allows the plant to use JA to fine-tune responses to multiple stresses^35^. The exaggerated root shortening and enhanced *A. brassicicola* susceptibility phenotypes of the *llp1-1, llp3,* and *RNAi:LLP1-3* plants suggest that there is a misregulation in signalling at a point upstream of one of JA’s two key pathway regulators, MYC2 and ERF1^36^. Transcript levels of the ERF1 pathway marker gene *PDF1.2* were reduced in MeJA-treated *RNAi:LLP1-3* plants, whereas the MYC2 pathway marker gene *VSP2* was not misregulated in the same plants (Fig. 3 and Supplementary Fig. 6). *ERF1* is a key transcription factor activated in conjunction with ethylene signalling, which is implicated in defence against necrotrophic pathogens^37,38^, and is strongly induced in response to salt stress^39^. The susceptibility of *llp1-1, llp3,* and *RNAi:LLP1-3* plants to the pathogen *A. brassisicola* thus further supports a misregulation of the ERF1-regulated branch of JA signalling in these mutants. Together, the data suggest that LLP1, 2, and/or 3 influence JA responses through ERF1 (Fig. 4C).

Although observed physiologically in all genotypes tested, misregulated SA-JA cross talk events were observed at the molecular level, *i.e. PDF1.2* and *PR1* transcript accumulation changes, only in *RNAi:LLP1-3* plants. This hints at possible additive roles of *LLP1, LLP2,* and *LLP3* in this process. During SAR, however, the roles of *LLP1* and *LLP3* appear to be spatially separated with *LLP1* acting systemically and *LLP3* promoting local SAR signal generation. This might explain why MeJA enhanced the systemic resistance of *llp1-1,* but not *RNAi:LLP1-3* plants against *Pst* if *LLP3* is required locally to drive JA-associated SAR signal generation or transmission through ERF1 (Fig. 4C). During SAR, SA-JA antagonism is observed locally, but not in the systemic tissues1^9,20^. Perhaps, *LLP1* fine-tunes incoming signals to avoid antagonistic trade-offs between SA- and JA-mediated defences in the systemic tissue during SAR (Fig. 4C).

The recently suggested spatial role of JA signalling in the perimeter of SA-induced HR lesions^21^ might explain how JA locally influences SAR signal generation. High SA levels in the core of the lesion promote *LLP1* transcript accumulation, while JA accumulation in the rim of the lesion drives up *LLP3* expression. We hypothesize that this signal is then relayed through the *ERF1* pathway affecting salt tolerance, defence against necrotrophic pathogens, and also driving SAR signal emission from this site (Fig. 4C). The role of *LLP1* allows this pathway to act synergistically with both the SA- and Pip-dependent systemic defence signals, creating an interwoven network necessary for SAR-associated defence priming. Priming of SA defences in the absence of deleterious effects on JA defences further assigns a high potential to LLP-associated SAR signalling components for application in future durable plant protection strategies. A possible exploitation of LLP-associated signaling moieties towards resource-efficient defence priming will be subject of further study.

## Methods

### Plant materials and growth conditions

*A. thaliana* ecotype Columbia-0 (Col-0) was used as the wild type control throughout all experiments. Transgenic lines *llp1-1, eds1-2, ald1, ggpps12, tps24-1, tps24-2,* and *RNAi:LLP1-3* have been described previously^5,6,34,40,41^. *RNAi_LLP1-3* line C3 13-1^6^ was used for all experiments. SALK_030762 with a T-DNA insertion in *LLP3* (At3G15356) was obtained from the Nottingham Arabidopsis Stock Center^42^, and propagated to homozygosity. Plants that were homozygous for the T-DNA insertion were used for all experiments and as the parental line for generating *llp3-LLP3:LLP3* complementation lines 3.02 and 4.01. For the latter, *LLP3:LLP3* constructs were generated from Col-0 wild type genomic DNA. The native promoter was chosen from ~2 kilo base pairs upstream to the *LLP3* transcriptional start site, and the *LLP3:LLP3* target sequence was isolated by PCR using the primers LLP3:LLP3-F and LLP3:LLP3-R (Supplementary Table 1). The resulting DNA fragment was cloned into pENTR™/ D-TOPO® (Invitrogen) and sequenced. The resulting construct was transferred to the binary Gateway® cloning vector pBGWFS7,0^43^, with the *GUS* sequence removed using the restriction enzyme NruI (pBGWFS7,0ΔGUS). The resulting binary vector was transformed into *Agrobacterium tumefaciens* strain GV3101 and used for plant transformation by floral dip^44^. Transgenic T1 plants were selected via 200g/L BASTA spray (Hoechst, Germany). Experiments were performed in T3 plants. *LLP3* transcript levels were determined by RT-qPCR as described below with the *LLP3* primer sets c1 or c2 (Supplementary Table 1).

Plants were grown on potting soil (without fertilizer) mixed with sand in 5:1 ratio, and 2 1 kept under short day conditions (10 hours (h) light with an intensity of 100 μE m^−2^ s^−1^ at 22°C and 14 h dark at 18°C, 70% relative humidity).

### Phytohormone treatments

To analyse *LLP1-3* transcript accumulation in response to phytohormone treatment, green tissues of 2- to 3-week old plants were sprayed until drop-off with 1 mM SA (Sigma Aldrich), 100 μM MeJA (Sigma Aldrich), or 100 μM ABA (Sigma Aldrich) dissolved in 0.1% MgCl_2_, 0.01% Tween® 20, and 0.025% MeOH. Plants of the same age were sprayed with 0.1% MgCl_2_, 0.01% Tween® 20, 0.025% MeOH as the mock control treatment. Leaf samples were taken at 8 and 24 h after treatment and flash frozen in liquid N2.

### Pathogen infection assays

*Pseudomonas syringae* pathovar *tomato* (*Pst*) and *Pst AvrRpm1* were maintained as described^5^. To induce a SAR response, plants were infiltrated in their first two true leaves with 1×10^6^ CFU/mL of *Pst/AvrRPM1.* Three days later, two systemic leaves were infiltrated with 1×10^5^ CFU/mL of *Pst*. The resulting *in planta* bacterial titres were determined at 4 dpi as described^6^. *A. brassicicola* was maintained on malt medium (3% malt extract (Merck), 1.5% agar-agar (Roth)) and transferred to oat plates (oats (Alnatura) in 1.5% w/v agar-agar) before experiments. Mycelium was solved in MKP buffer (62mM KH_2_PO_4_, 0.01% glucose, 0.01% Tween® 20) until a concentration of 200 spores/μL was achieved. Plants were inoculated by placing 3 μL droplets onto the third and fourth true leaf. The resulting lesion sizes were determined at 5 dpi using ImageJ. Cell death was visualized using trypan blue staining as described^45^.

### Petiole exudate experiments

Petiole exudate experiments were performed as described^6^. In short, *Pst AvrRpm1*-inoculated leaves were cut in the middle of the rosette at 24 h post-inoculation, and incubated with their petioles in 1 mM EDTA. After 1 h 6 leaves per exudate were transferred to 2.0 mL of sterilized water and allowed to exude for 48 h. The resulting PEX solutions were filter-sterilized (Millipore, 0.22 μm) and supplemented with MgCl_2_ to a final concentration of 1 mM. 24 h after syringe infiltration of the PEX in leaves of naïve recipient plants, the infiltrated leaves were inoculated with 10^5^ cfu mL^−1^ of *Pst*, *in planta* titres of which were determined at 4 dpi as described above.

### Root length inhibition assays

For root growth inhibition measurements, seedlings were sterilised in 75% followed by 100% EtOH (Merck), dried, and sown on 1x Murashige Skoog medium including vitamins (Duchefa) with 0.1% cefotaxim (Acros Organics) and 0.25% Carbenicillin (Roth). Seedlings were transferred after 6 days to treatment plates containing either 10 μM ABA, 100 mM NaCl, or 40 μM MeJA (Sigma-Aldrich), or to control MS plates. All plates were placed upright in the growth chamber under long day conditions, and the seedlings were photographed 6 and 12 days post-transfer. Root length was measured using ImageJ. The seedlings were harvested, pooled per genotype and treatment, and flash frozen in N_2_ for RNA extraction.

### Phytohormone content measurements

ABA in seedlings was measured as described^46^. In short, the frozen material was spiked with 10 ng ABA-d_6_ and incubated with 40% acetonitrile (ACN) for 30 min prior acidification with phosphoric acid and extraction with *tert-butyl* methyl ether. The organic extract was passed over a Chromabond NH2 500 mg solid phase extraction column (Macherey-Nagel, Düren, Germany). The eluate was diluted with distilled water and passed over a Chromabond C18ec 100 mg solid phase extraction column (Macherey-Nagel). The eluate was evaporated in a vacuum concentrator, dissolved and fractionated by RP-HPLC using a Nucleodur 100-5 C18ec 125×4.6 mm column (Macherey-Nagel). The ABA-containing fraction was collected, evaporated to dryness and methylated with 2 M (trimethylsilyl)diazomethane/methanol = 1:19. After evaporation, the residue was dissolved in ACN and analysed by gas chromatography-mass spectrometry using a VF-5ms column (Agilent, St. Louis, MO, USA) and helium as carrier gas at a flow rate of 1.5 ml/min. The ions with *m/z* 190 (ABA) and 194 (ABA-d6) were used for quantification and the ions at 134 and 162 (ABA) and 138 and 166 (ABA-d6) were used as qualifiers.

The phytohormones ABA, SA and JA were measured in mature plants according to^46^ using a versatile UHPLC-MS/MSMRM system. The plant material (50-250 mg) was placed in 2 mL bead beater tubes (CKMix-2 mL, Bertin Technologies, Montigny-le-Bretonneux, France). An aliquot of the internal standard (20 *μ*L), containing ABA-d_6_ (2.5 *μ*g/mL), SA-d_4_ (2.5 *μ*g/mL), and JA-d_5_ (25 *μ*g/mL) in acetonitrile was added to the plants and incubated for 30 min at room temperature. After extractive grinding with ethyl acetate (1 mL) in a bead beater (Precellys Homogenizer, Bertin Technologies, Montigny-le-Bretonneux, France) the supernatant was membrane filtered (0.45 *μ*m), evaporated to dryness, resolved in acetonitrile (70 *μ*L) and injected into the LC-MS/MS-system (2 *μ*L).

For LC-MS/MS analysis a QTRAP 6500^+^ mass spectrometer (Sciex, Darmstadt, Germany) was used to acquire electrospray ionization (ESI) mass spectra and product ion spectra. Negative and positive ions were detected in the scheduled multiple reaction monitoring (MRM) mode.

For analysis of ABA, SA and JA, the MS/MS parameters were tuned to achieve fragmentation of the [M-H]^−^ and [M+H]^+^ molecular ions into specific product ions to receive a qualifier and a quantifier transition for every compound.

Chromatography was performed by means of an ExionLC UHPLC system (Shimadzu Europa GmbH, Duisburg, Germany) equipped with a Kinetex F5 column (100 × 2.1 mm, 100 Å, 1.7 *μm,* Phenomenex, Aschaffenburg, Germany). Operated with a flow rate of 0.4 mL/min using 0.1% formic acid in water (v/v) as solvent A and 0.1% formic acid in acetonitrile (v/v) as solvent B, chromatography was performed with the following gradient: 0% B held for 2 min, increased in 1 min to 30% B, in 12 min to 30% B, increased in 0.5 min to 100% B, held 2 min isocratically at 100% B, decreased in 0.5 min to 0% B, held 3 min at 0% B. Data acquisition and instrumental control were performed using Analyst 1.6.3 software (Sciex, Darmstadt, Germany).

### Salt pouring experiments

Plants were watered with distilled water or 300 mM NaCl three times with four day intervals starting from 4 weeks after germination. Leaf tissue was harvested 4 days after the final salt treatment, weighed, and flash frozen in liquid N_2_.

### RNA isolation and RT-qPCR

Total RNA was extracted from leaves and seedlings using TriReagent (Sigma-Aldrich) following the manufacturer’s instructions. cDNA was synthesized using SuperScriptII reverse transcriptase (Invitrogen). Real-time quantitative PCR was performed using the Sensimix SYBR low-rox kit (Bioline) and the primers in Supplementary Table 2, with *UBIQUITIN* as the reference gene. Endogenous *LLP3* transcript accumulation was determined with the primers LLP3-F and LLP3-R. qPCR was performed on a 7500 real-time PCR system (Applied Biosystems). Transcript accumulation was analysed using the 7500 Fast System Software 1.3.1.

### Drought assay

The progressive drought experiment was performed as described^47^. In brief, Arabidopsis plantlets were exposed to a slowly increasing water deficit by minimizing evaporation and withholding watering under short day conditions (8 h light). Water consumption per plant was recorded from 18 to 73 days after seeding. Above-ground material was used for determining the dry-weight biomass and WUE was expressed as the ratio of biomass to consumed water.

### Statistics

Data was analysed in GraphPad Prism 8 for Windows. If necessary, outliers were removed using a Grubbs’ test (α=0.05). Normal distribution of the data was checked using D’Agostino Pearson (α=0.01). Data that showed normal distribution was tested for significance using an unpaired one-way ANOVA with Tukey’s multiple comparison test, and data that was not normally distributed was tested using a Kruskal-Wallis test with a Dunn’s multiple comparison test.

## Supplementary Material

**Supplementary Table 1** Primers for LLP3:LLP3 construct generation and qPCR

**Supplementary Table 2** Primers for qPCR

**Supplementary Figure 1** BTH induces transcript accumulation of *LLP1.*

**Supplementary Figure 2** *LLP3* transcript levels in *llp3* and *llp3-LLP3:LLP3* complementation lines

**Supplementary Figure 3** *LLP1* moderately influences primary root growth.

**Supplementary Figure 4** *llp1-1* and *RNAi:LLP1-3* lines do not show an altered response to ABA

**Supplementary Figure 5** *LLP1-3* do not affect the response to drought stress.

**Supplementary Figure 6** JA-associated marker gene expression

**Supplementary Figure 7** LLP1-3 do not influence phytohormone accumulation in response to salt

**Supplementary Figure 8** Salt- and MeJA-induced primary root growth inhibition in seedlings of SAR-associated mutant lines

## Acknowledgments

We thank Miriam Lenk for critically reading the manuscript. This work was funded by the Deutsche Forschungsgemeinschaft (DFG) as part of SFB924 (projects A12 (BP), B01 (EG), B12 (CD), and project B06 to ACV).

