## Supplementary Tables and Figures for "Local jasmonic acid cues drive systemic acquired resistance signal generation"

**Supplementary Table 1** Primers for *LLP3:LLP3* construct generation (upper panel) and qPCR (lower panel)

| Primer name | Sequence (5’ → 3’) |
|---|---|
| **LLP3:LLP3-F** | CACCAGCAATCATCAAAAGACACAAGGAA |
| **LLP3:LLP3-R** | TTAGTTTTCAAACGACCAGCTCCA |

| Primer name | Sequence (5’ → 3’) |
|---|---|
| **C1-LLP3-F** | CCATTACCATCGCCCCTGAA |
| **C1-LLP3-R** | CCCATGGAACCAGCAAAACC |
| **C2-LLP3-F** | ACCGAAGAGGCCTTTGATCC |
| **C2-LLP3-R** | ACCAGCAAAACCAGCGTACA |

**Supplementary Table 2** Primers for qPCR

| Primer name | Sequence (5' to 3') |
|---|---|
| **LLP1-F** | TGAGTAAACAGCAGTTACGA |
| **LLP1-R** | TGACGCCATCAGAAGCAGGA |
| **LLP2-F** | CCCGAGACGAGAACCCAT |
| **LLP2-R** | GGAGTTATCAGCTTCCGGGG |
| **LLP3-F** | TTTGGAGCTGGTCGTTTG |
| **LLP3-R** | ATTCACTCTACAACAATT |
| **PR1-F** | CTACGCAGAACAACTAAGAGGCAAC |
| **PR1-R** | TTGGCACATCCGAGTCTCACTG |
| **RAB18-F** | TTCGGTCGTTGTATTGTGCTTT |
| **RAB18-R** | CCAGATGCTCATTACACACTCATG |
| **PDF1.2-F** | CCAAGTGGGACATGGTCAG |
| **PDF1.2-R** | ACTTGTGTGCTGGGAAGACA |
| **VSP2-F** | GTTAGGGACCGGAGCATCAA |
| **VSP2-R** | AACGGTCACTGAGTATGATGGGT |

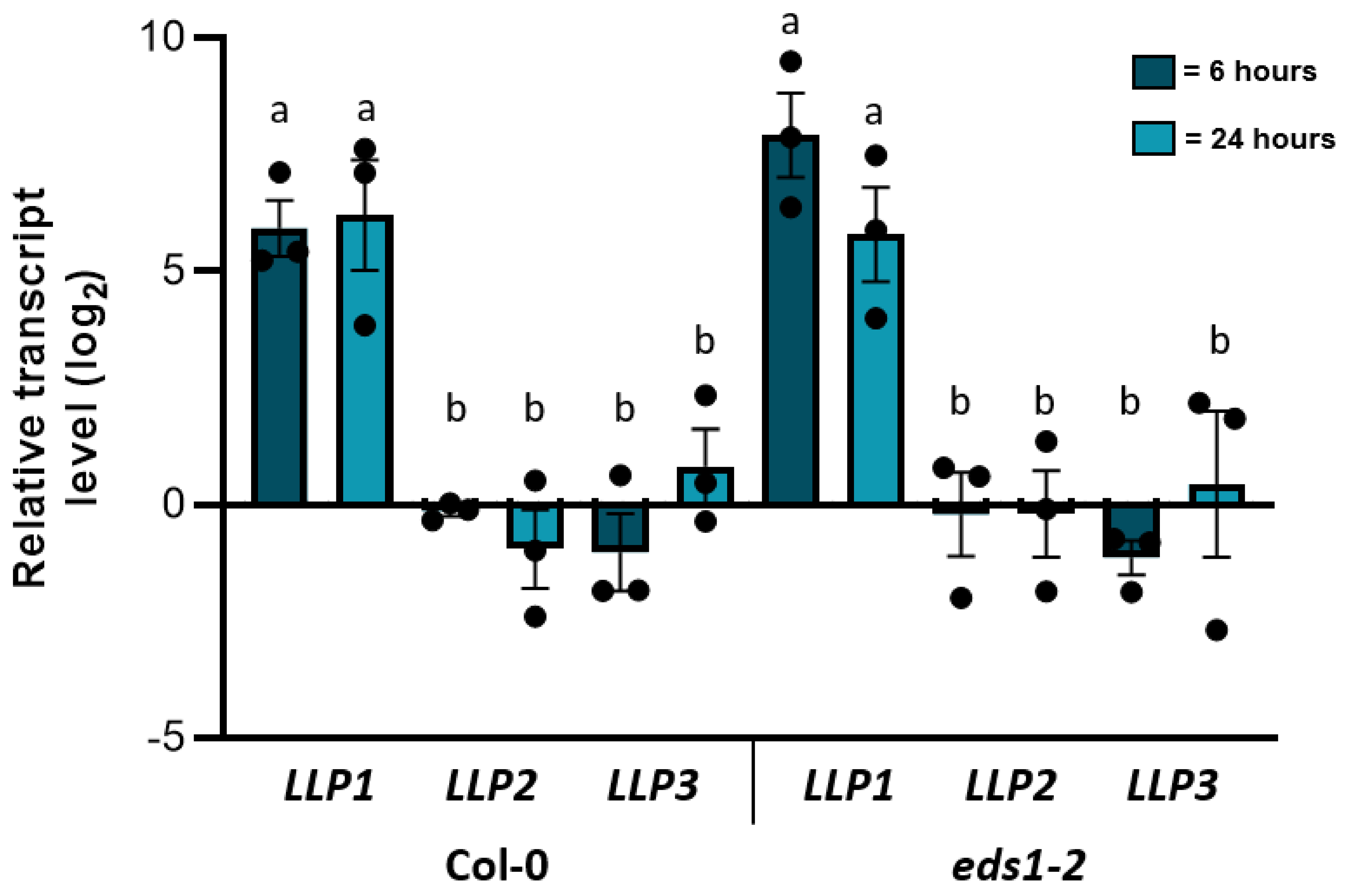


**Supplementary Figure 1** BTH induces transcript accumulation of *LLP1*. Transcript accumulation of *LLP1, LLP2,* and *LLP3* normalized to *UBIQUITIN* in 3-week-old Col-0 and *eds1-2* after spray-treatment with 1 mM BTH. Transcript accumulation was determined at 6 and 24h after treatment by RT-qPCR, and is shown relative to that in the appropriate mock controls. Dots indicate individual results from biologically independent experiments. Bars represent the $log_2$(mean) ± SEM. The letters above the bars indicate statistically significant differences (one-way ANOVA , n=3, P= ≤0.05,  for Col-0 6h F=17.12, DF=53, for Col-0 24h F=15.33, DF=53, for *eds1-2* 6 h F=11.33, DF=53, for *eds1-2* 24h F=10.39, DF=53).

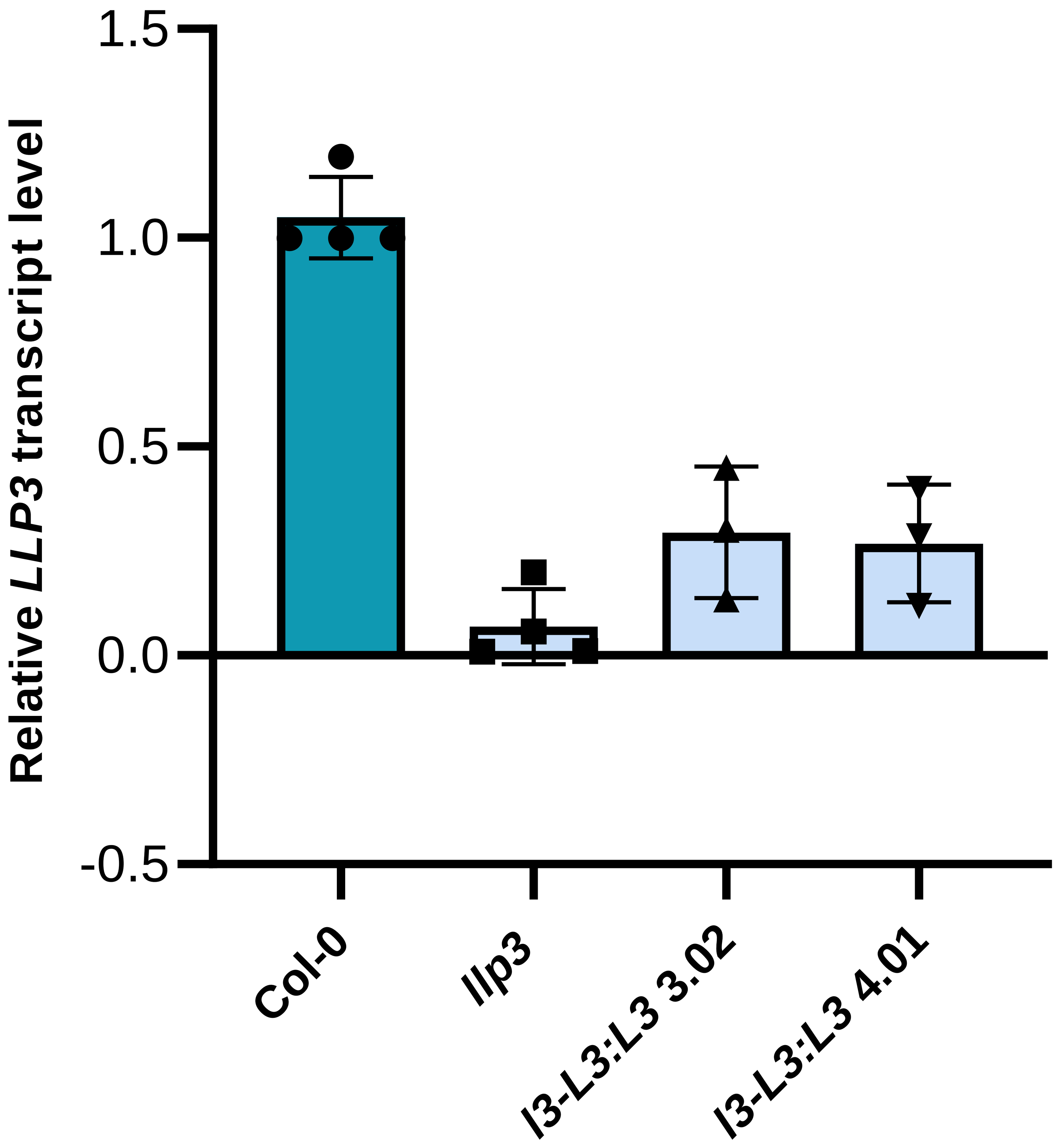


**Supplementary Figure 2** *LLP3* transcript levels in *llp3* and *llp3-LLP3:LLP3* complementation lines 3.02 and 4.01. *LLP3* transcript accumulation was determined in 5-to-6 week-old plants and normalized to that of *UBIQUITIN*. *l3-L3:L3* 3.02 and *l3-L3:L3* 4.01 are two biologically independent transgenic lines carrying the *LLP3:LLP3* transgene driving ectopic *LLP3* expression from its native promoter in the *llp3* mutant background. Bars represent the average of the indicated results ± SEM (one-way ANOVA, n=4 for Col-0 and *llp3,* n=3 for *l3-L3:L3 3.02* and *l3-L3:L3 4.01,* F=51.41, DF=13).

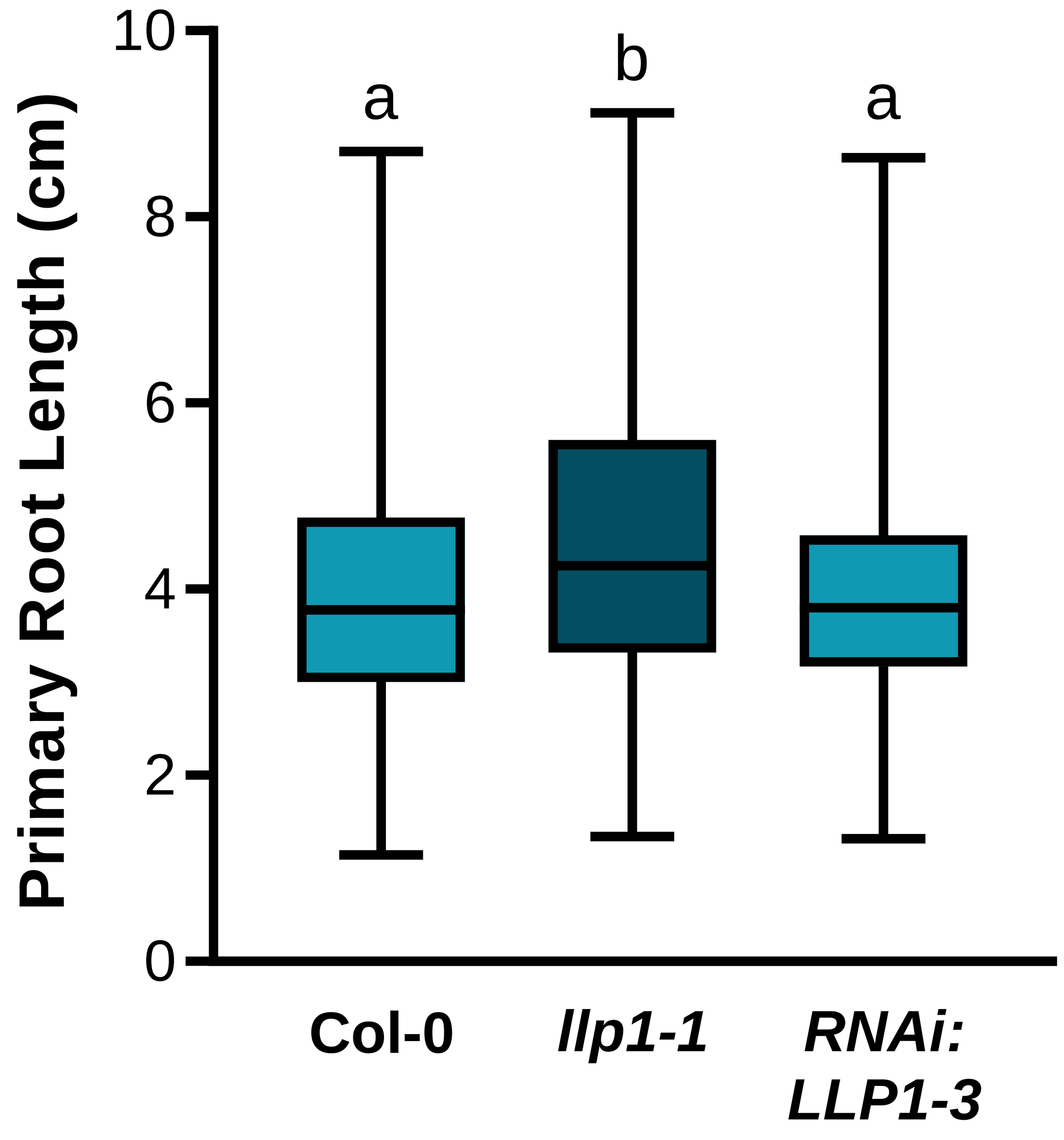


**Supplementary Figure 3** *LLP1* moderately influences primary root growth. 6 day old seedlings were transferred from germination plates to fresh control plates (in parallel with the experimental treatments shown in Figs. 2 and 3). The primary root length of wild type, *llp1-1* and *RNAi:LLP1-3* was measured 12 days after transfer. Box plots indicate the average root length ± min and max values. The letters above the box plots indicate statistically significant differences (Kruskal-Wallis test, P=<0.05, KW test statistic=14.41, Col-0 n=447, *llp1-1* n=200, *RNAiLLP1-3* n=239 from 9 biologically independent experiments).

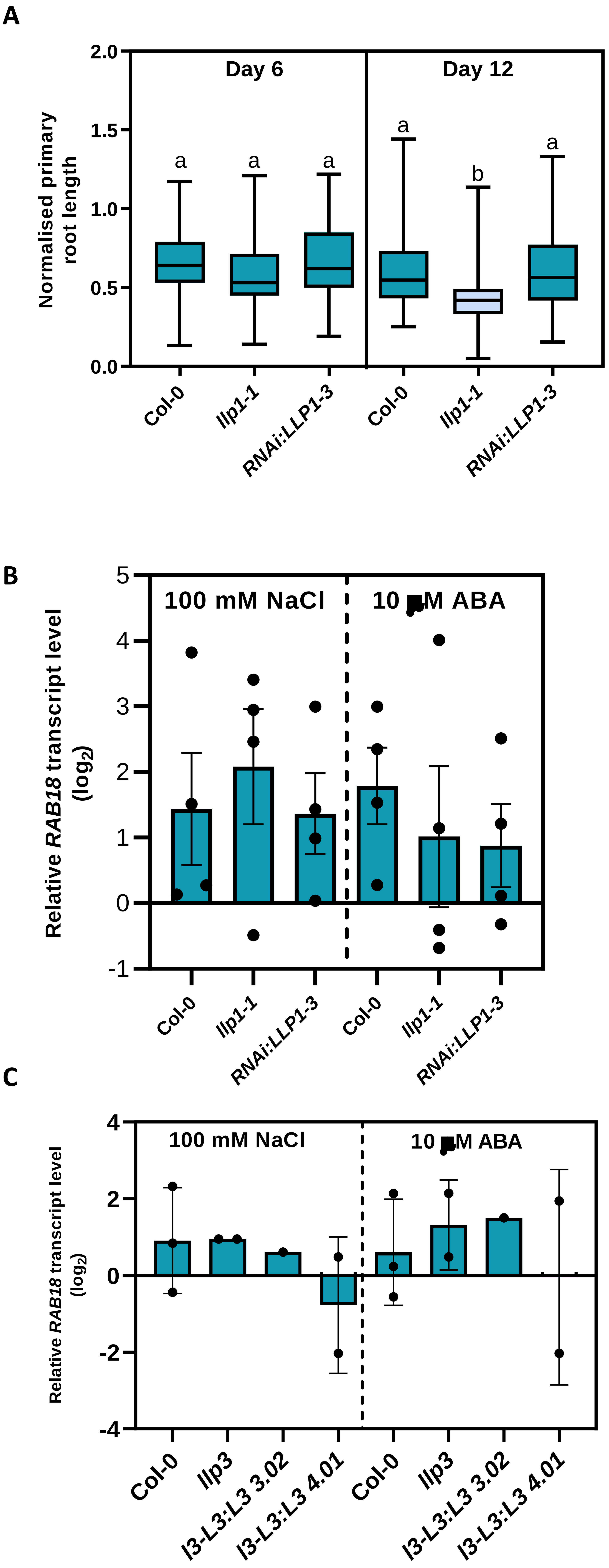


**Supplementary Figure 4** *llp1-1* and *RNAi:LLP1-3* lines do not show an altered response to ABA treatment. 6 day old seedlings were transferred to treatment plates containing 100 mM NaCl or 10 µM ABA. (A) Primary root growth of Col-0 wild type, *llp1-1* and *RNAi:LLP1-3* on treatment plates containing ABA was determined at 6 and 12 days after transfer and normalized to that on control plates. Box plots indicate average normalised root length ± min and max values. The letters above the box plots indicate statistically significant differences. (Kruskal-Wallis test, P=<0.05, KW test statistic=125.4, for day 6 Col-0 n=321 , llp1-1 n=136, RNAi:LLP1-3 n=125, for day 12 Col-0=290, llp1-1 n=105, RNAiLLP1-3 n=138). Data were merged from 4 biologically independent replicates, and exaggerated ABA-induced root growth inhibition in *llp1-1* was significant in only 1 replicate. (B/C) Salt- and ABA-induced *RAB18* transcript levels were similar in all genotypes. Transcript levels of *RAB18* were normalized to that of *UBIQUITIN* and are shown relative to the normalized *RAB18* transcript levels in the appropriate mock controls. Dots indicate data from biologically independent experiments. Bars represent the log$_2$(mean) ± SEM (B: NaCl: one-way ANOVA, n=4, P=<0.05, F=0.9399, DF=35, ABA: one-way ANOVA, n=4, P=<0.05, F=0.7500, DF=35; C: NaCl: one-way ANOVA, Col-0 n=3, *llp3* n=2, *l3-L3:L3 3.02* n=1, *l3-L3:L3 4.01* n=2, P=<0.05, F=0.4480, DF=7, ABA: one-way ANOVA Col-0 n=3, *llp3* n=2, *l3-L3:L3 3.02* n=1, *l3-L3:L3 4.01* n=2, P=<0.05, F=0.0878, DF=7).

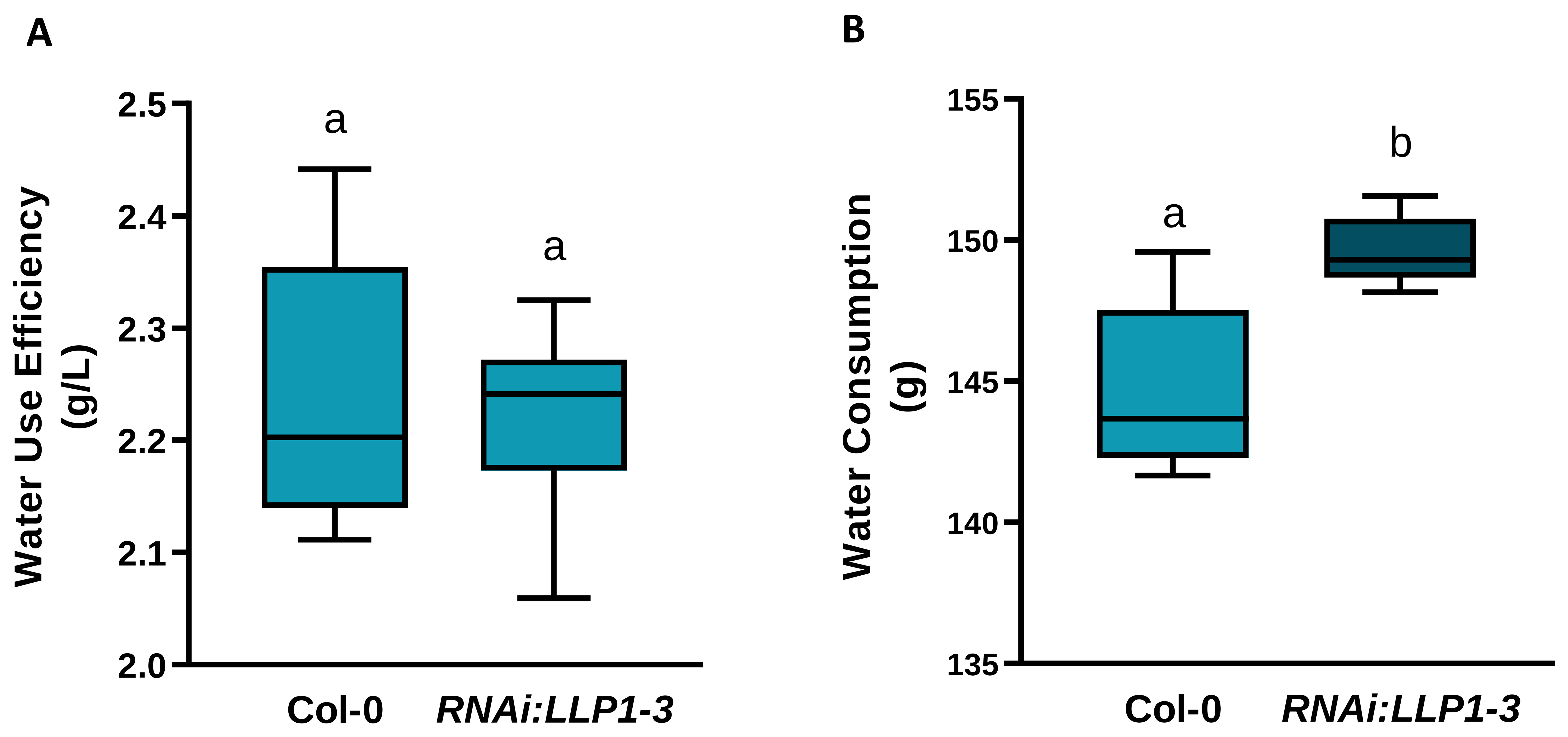


**Supplementary Figure 5** *LLP1-3* do not affect the response to drought stress. Col-0 wild type and *RNAi:LLP1-3* plants were well-watered for 21 days, and then placed under progressive drought conditions. (A) Water use efficiency is the same in both genotypes. Box plots indicate the average water use efficiency ± min and max values (n=12, unpaired two-tailed t test, t=0.3456, df=21). (B) Mean water consumption under drought conditions up to 56 days. Box plots indicate the mean water consumption ± min and max values. The letters above the box plots indicate statistically significant differences (n=12, unpaired two-tailed t test, t=5.487, df=21). Although there is a statistical difference, this is not considered a notable physiological difference when compared to ABA-related drought mutants.

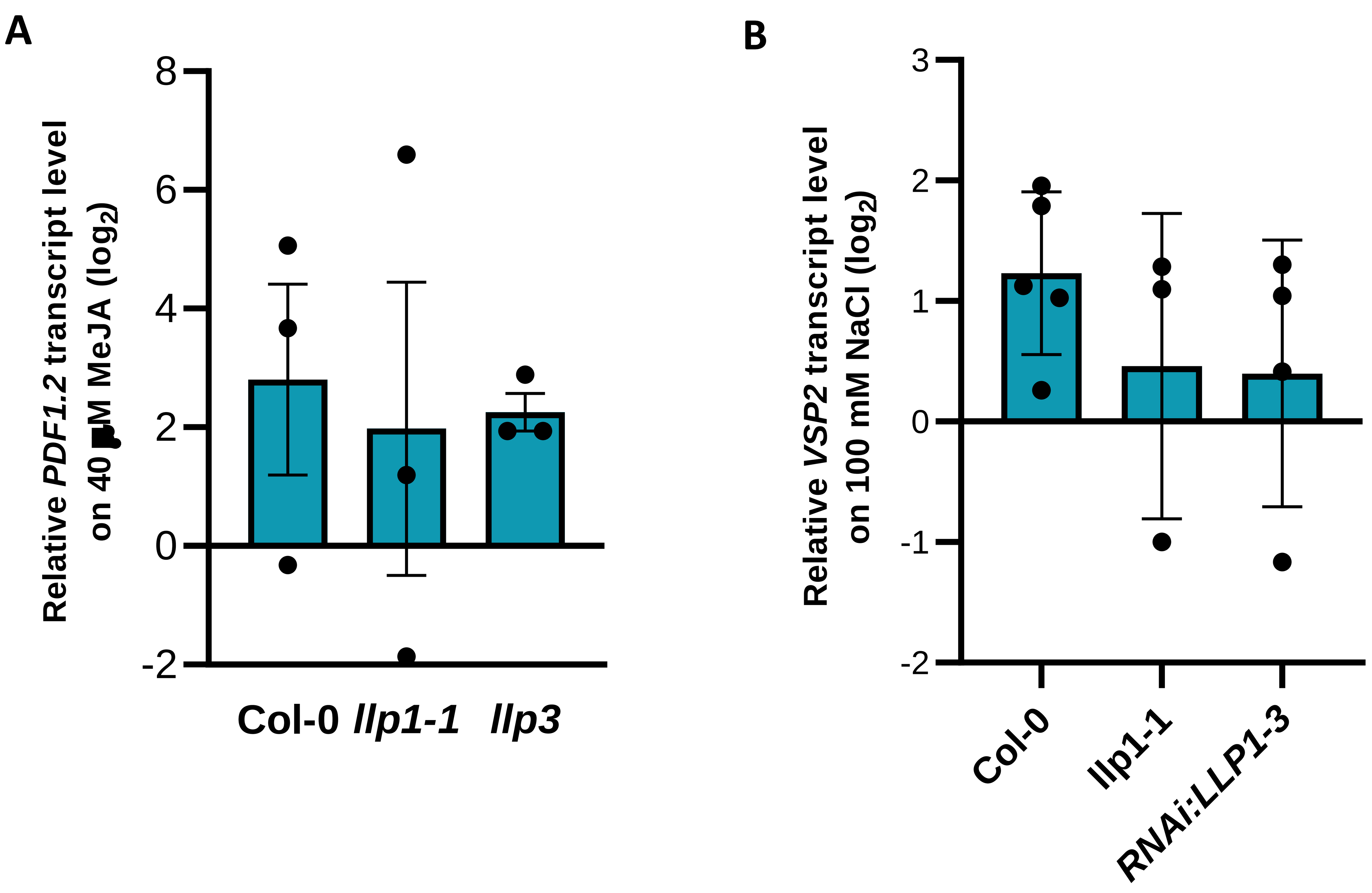


**Supplementary Figure 6** JA-associated marker gene expression. (A) MeJA-induced *PDF1.2* transcript accumulation was not different in *llp1-1* or *llp3* as compared to that in Col-0 wild type seedlings. Seedlings of the indicated genotypes were germinated on control MS media, and after 6 days transferred to either further control plates, or to MS plates supplemented with 40 µM MeJA. *PDF1.2* transcript levels were determined by qRT-PCR 12 days after transfer and normalised to that of *UBIQUITIN*. Normalized transcript levels are shown relative to those in the appropriate controls. Dots indicate data from biologically independent experiments. Bars represent the $\log_2$(mean) ± SEM (One-way ANOVA, n=3, P=<0.05, F=0.1166, DF=10). (B) *VSP2* transcript levels on 100 mM NaCl are the same in *llp1-1* or *llp3* as compared to those in Col-0 wild type seedlings. Seedlings were treated as described in (A). *VSP2* transcript levels were determined by qRT-PCR 12 days after transfer and normalised to that of *UBIQUITIN*. Normalized transcript levels are shown relative to those in the appropriate controls. Dots indicate data from biologically independent experiments. Bars represent the $\log_2$(mean) ± SEM (One-way ANOVA, P=<0.05, F=1.883, DF=20, Col-0 n=5, *llp1-1* n=3, C3 13-1 n=4).

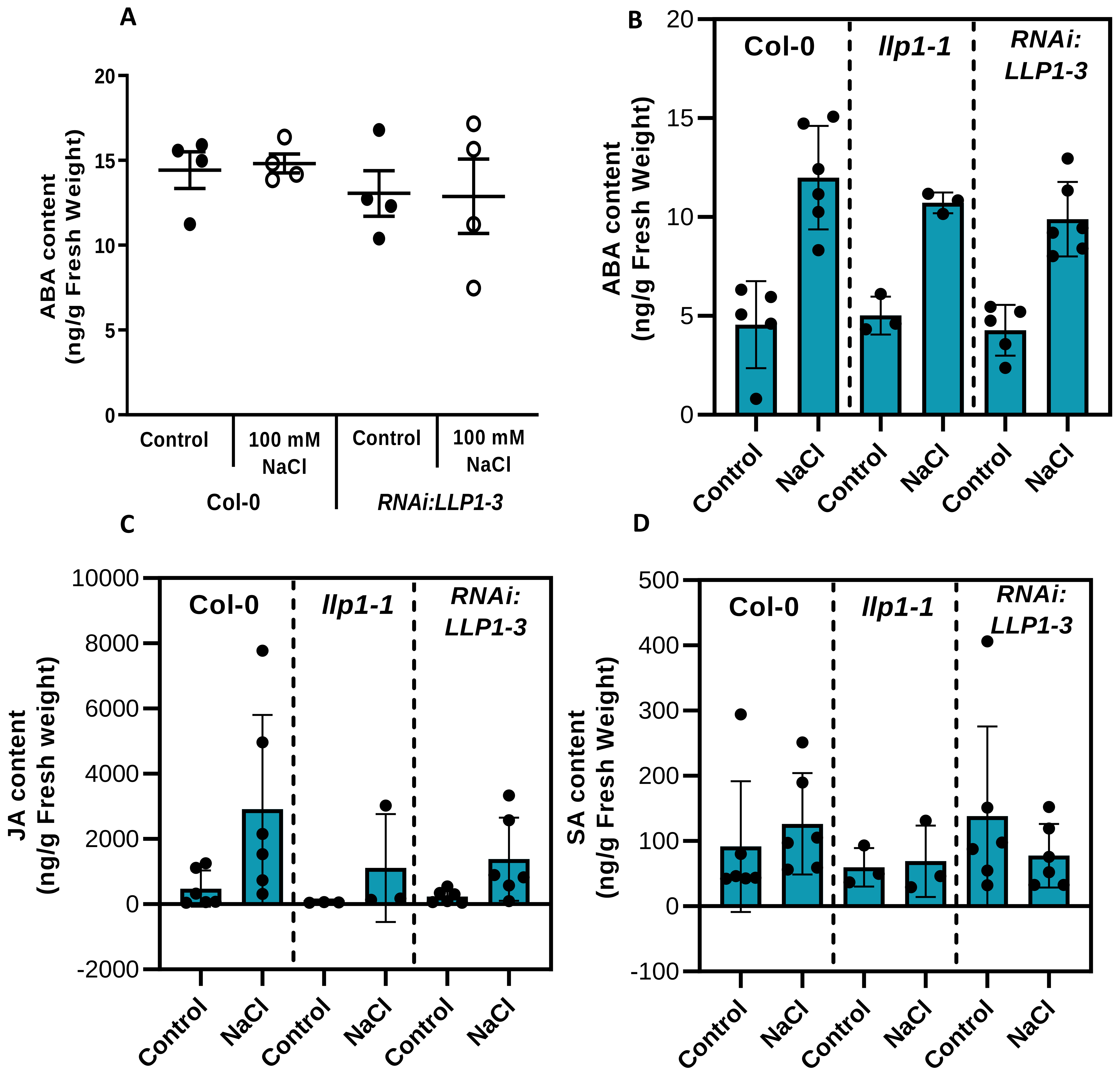


**Supplementary Figure 7** LLP1-3 do not influence phytohormone accumulation in response to salt. (A) ABA content 12 days after transfer of seedlings from control to further control or treatment MS plates supplemented with 100 mM NaCl. Individual data points are derived from biologically independent experiments. Shown are average ± SEM (one-way ANOVA, n=6, P=<0.05, F=0.4622, DF=15). (B-D) Phytohormone content in 5-week-old plants after 12 days of exposure to 300 mM NaCl by irrigation. Individual data points are derived from biologically independent experiments. (B) ABA content. Bars indicate average ± SEM (one-way ANOVA, P=<0.05, , for Col-0 and *RNAiLLP1-3* n=6, *llp1-1* n=3. , F=0.1859 DF=12 ). (C) JA content. Bars indicate average ± SEM (Kruskal-Wallis test, , for Col-0 and *RNAiLLP1-3* n=6, *llp1-1* n=3, KW=0.1403). (D) SA content. Bars indicate average ± SEM (Kruskal-Wallis test, , for Col-0 and *RNAiLLP1-3* n=6, *llp1-1* n=3, KW=0.302).

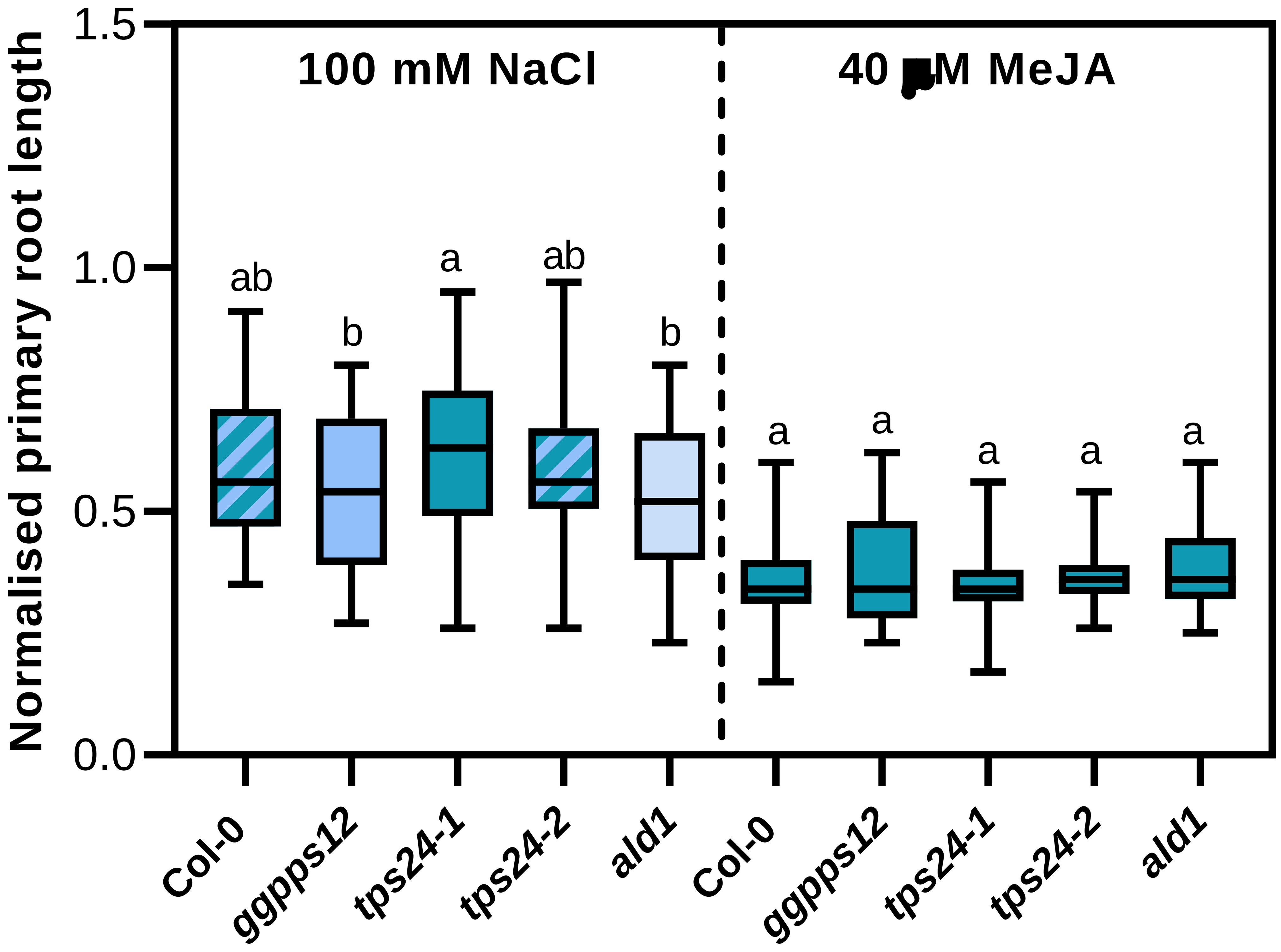


**Supplementary Figure 8** Salt- and MeJA-induced primary root growth inhibition in seedlings of SAR-associated mutant lines. Tested genotypes include: pipecolic acid-deficient *ald1* and monoterpene-compromised *geranyl geranyl diphosphate synthase12* (*ggpps12*) and *terpene synthase 24* (*tps24*) mutants. The seedlings were germinated on control plates and transferred to further control or treatment plates after 6 days. Roots were measured after 12 days of treatment with either 100 mM NaCl or 40 µM MeJA. Values shown are primary root length on treatment plates normalised to primary root length on control plates. Box plots indicate average normalized root length ± SEM. The letters above the box plots indicate statistically significant differences (Kruskal-Wallis test, P=<0.05,Col-0 n=87, n=103. *ggpps12 208* n=117, n=117. *tps12 055* n=80, n=117. *tps12 127* n=85, n=118. *ald1* n=110, n=116 from 2 biologically independent experiments. For 100mM NaCL, KW statistic=25.00, for MeJA, KW statistic=8.924)
